# Beyond Expression Prediction: Benchmarking Differential Expression Classification in Single-Cell Perturbation Models

**DOI:** 10.64898/2026.07.20.739620

**Authors:** Junwei Sun, Yuxun He, Ouyang Zhu, Yiqun T. Chen

## Abstract

Accurate predictions of transcriptomic responses to genetic perturbations could unlock our understanding of gene functions and regulatory networks. While a growing number of methods and benchmarks target this task, existing evaluations focus on mean expression accuracy alone. This overlooks differential expression (DE), which captures both mean and variance and forms the basis for biological interpretation and experimental follow-up. Here, we systematically evaluate a diverse set of deep learning and non-deep-learning methods for their ability to predict DE outcomes under two generalization regimes: unseen perturbations within the same cell line, and unseen cellular contexts across cell lines. We find that simple baselines, such as embedding-based nearest neighbors, are competitive and often outperform specialized deep learning models for DE classification across datasets and evaluation metrics. We further show that sparsity calibration, motivated by the structure of single-cell data, substantially improves DE classification for deep learning models that do not explicitly account for sparsity. Together, our findings establish practical baselines and evaluation principles for benchmarking perturbation models on DE prediction.

## 1 Introduction

Understanding how cells respond to external perturbations such as small molecules and genetic modifications could unlock much-needed understanding of gene regulation and cellular biology. Recent technologies, such as Perturb-seq [1], pair CRISPR-based perturbations with single-cell RNA sequencing to measure transcriptomic profiles after gene knockdowns at scale. Perturb-seq experiments have yielded meaningful biological insights: for instance, perturbing *ANK2*, a gene linked to neurodevelopment, was found to significantly upregulate an interneuron-specific gene module, suggesting a role in interneuron maturation beyond its known functions [2].

Despite their potential to further biological understanding, perturbational screens remain expensive. As a result, computational models that predict transcriptomic responses to genetic perturbations have become a major focus in computational biology and artificial intelligence, with the hope that accurate predictions could inform or even substitute for costly wet-lab experiments — an aspiration sometimes captured by the term “virtual cells.” [3, 4] A key downstream application of these models is differential expression (DE) analysis [5–7], which classifies each gene’s response to a perturbation as up-regulated, down-regulated, or unchanged based on whether its expression significantly increases, decreases, or remains stable relative to an unperturbed control, respectively.

Existing approaches [4] broadly fall into two categories. The first predicts continuous gene expression profiles, which are subsequently discretized into DE categories via statistical tests; representative methods include GEARS [8], scGPT [9], GenePert [10], VAE-based approaches (e.g., scVI [11], scGen [12]), transformer-based method (e.g., STATE [13]), and models utilizing large language model (LLM) derived embeddings such as Scouter [14] and scPert [15]. The second category directly models DE outcomes as a classification problem, often leveraging LLM reasoning. Notable examples include PerturbQA [16], SynthPert [17], and rbio1 [18]. Beyond these, agentic approaches such as VCHarness [19] and CellForge [20] autonomously construct perturbation-response models by combining AI coding agents with multimodal biological foundation models.

As the model space rapidly expands, rigorous and standardized evaluation has become increasingly important, which motivated a growing number of benchmarking effort [21–27]. Despite the extensive coverage of cell lines, models, and evaluation metrics of current benchmarks, DE classification has received little attention as an evaluation target. In particular, benchmarking model performance on more biologically interpretable DE labels (i.e., up-regulation, down-regulation, or no significant change) could be more relevant in downstream experimental design and validation, where a predicted effect’s significance must be judged against the variability inherent in the data rather than taken at face value [2, 28, 29]. In greater detail, existing benchmarks primarily assess models on continuous expression metrics (e.g., Pearson correlation, mean absolute error). For those that do evaluate DE prediction, the most common choice is a top-*k* DE gene overlap score: the overlap between the top-*k* predicted and top-*k* true DE genes (both ranked by true absolute log_2_ fold change), divided by *k*. This metric, however, overlooks class-wise performance across up-regulated, down-regulated, and unchanged genes. Furthermore, as emerging methods increasingly model discrete DE outcomes directly [16–18], systematically comparing simple discrete classifiers against DE predictions derived from more complex deep learning models could help establish best practices for DE prediction.

We therefore present a benchmarking study centered on discrete differential gene identification. We evaluate five deep learning models and three non-deep-learning methods across four cell-line datasets and two generalization regimes: unseen perturbations within the same cell line, and unseen cellular contexts. Non-deep-learning methods include a perturbation majority vote (i.e., predicts the DE outcome of a gene by its most common behavior under training perturbations), embedding-based k-nearest neighbor classifiers (i.e., classification based on KNN where perturbations are represented by embeddings, specifically we use the GenePT embeddings [30]), and a regularized logistic regression with perturbation embeddings as features. Deep learning methods include scVI [11], which we use both sampling-based and mean-expression predictions, referred to as scVI_sampling and scVI_mean respectively, Scouter [14], and a variational autoencoder adapted from scGen (referred to as VAE_scGen) for the within-cell-line setting [12]; STATE [13], TranScouter [31], scVI_sampling, scVI_mean, and VAE_scGen are additionally evaluated in the cross-cell-line setting (see Figure 1). For details on individual models, see Methods at section 3. We evaluate across four widely-used essential gene screen cell lines: K562 [32], RPE1 [32], Jurkat [33], and HepG2 [33]. We also explore the effect of distribution calibration and subset sampling on the benchmarking results (Figure 1). Our key findings are:

1. **Simple classification baselines are competitive with, and often out-perform, deep learning models on DE classification**, across both unseen-perturbation and unseen-cellular-context generalization tasks. Simple baselines achieve this by providing *high precision* predictions (i.e., more precise DE gene identification) while deep learning models tend to favor *higher recall* at the cost of precision (i.e., more false positive DE calls).
2. Model rankings depend on the choice of evaluation metric, and no single model uniformly outperforms across all settings. Moreover, **better performance on continuous expression metrics does not imply better DE prediction performance**, underscoring the need for a tailored metric panel when evaluating single-cell perturbation models.
3. **Calibrating predictions to the sparsity of real single-cell data consistently improves DE classification** for deep learning models that do not explicitly account for sparsity, while offering limited benefit to models with builtin distributional regularization. This points to calibration as a practical lever for improving existing methods.
4. **Data curation could inflate reported performance**. While subsampling rare classes could be an effective way to train models in the presence of rare DE classes, model performance and ranking on the subsampled data could differ drastically from performance on the full dataset, which is more practically relevant to real-world deployment conditions.

**Fig. 1:**
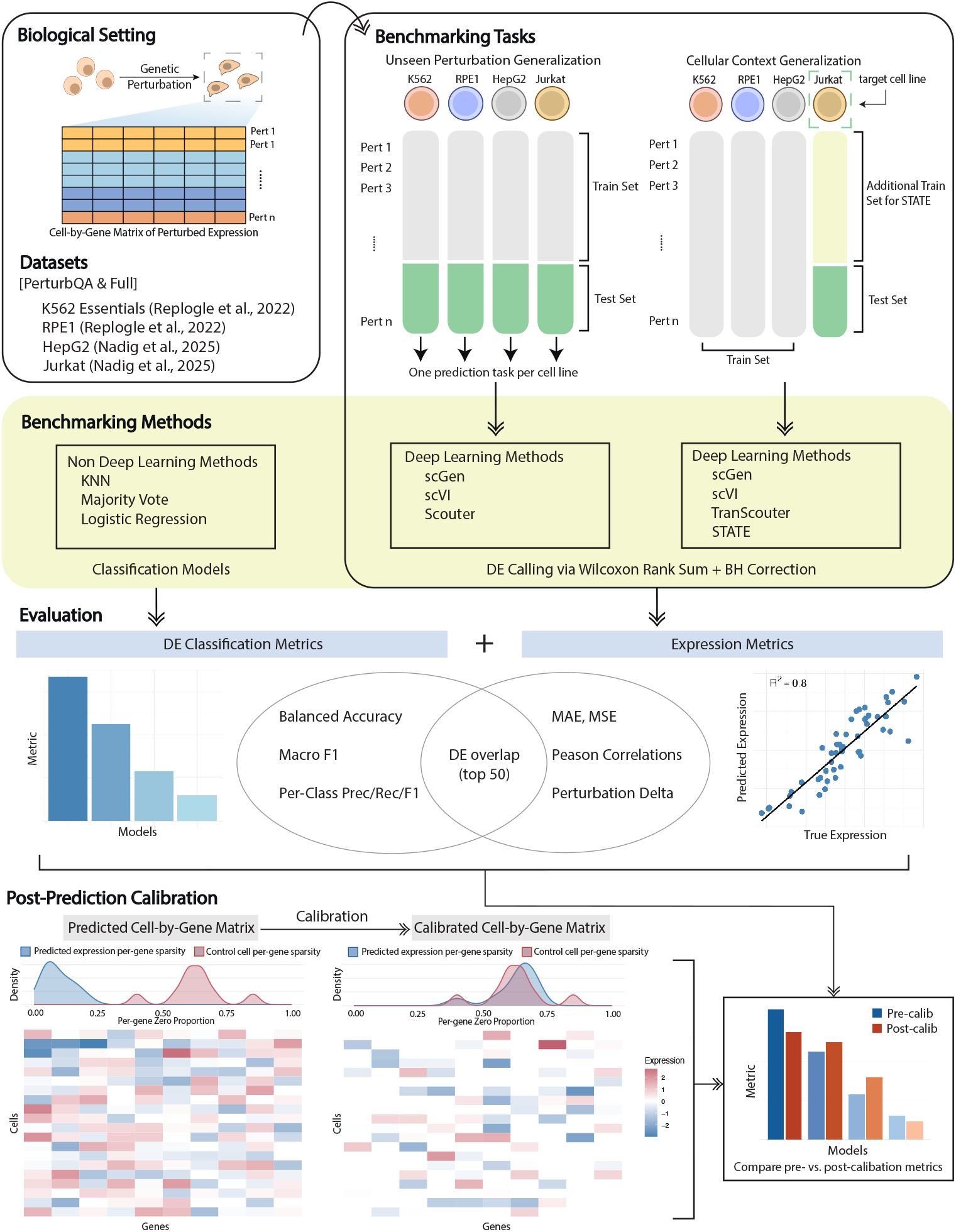
Study overview. **Biological Setting**: Single-cell perturbation data are collected through sequencing individual cells that have undergone genetic perturbations. **Datasets**: Four datasets corresponding to four cell lines collected this way are used for this study. Each dataset has a full, non-sampled version as well as a PerturbQA-downsampled variant that selects for highly significant DE pairs and downsamples non-DE pairs. **Benchmarking Tasks**: On both the full and PerturbQA-downsampled datasets, we evaluated two benchmarking tasks: unseen perturbation generalization and cellular context generalization. **Benchmarking Methods**: Each task is evaluated for three types of non-deep-learning methods and a varying number of deep learning methods. **Evaluation**: We used both DE classification and continuous expression metrics to report performance. **Post-Prediction Calibration**: Post-prediction, we further evaluated how adjusting for sparsity in the predicted expression space might impact model performance on DE classification.

### Related Work

Several recent work has focused on benchmarking single-cell perturbation models. Ahlmann-Eltze et al. [21] compared seven deep learning methods against simple linear baselines on dataset by Norman et al. [34], finding that no deep learning model outperforms simple methods on continuous expression metrics including root mean squared error and Pearson correlations. Wei et al. [24] extended this evaluation to 27 methods across 29 datasets using six complementary metrics, and Radig et al. [23] developed scArchon, a benchmarking platform evaluating models through a composite of statistical (t-tests, Wasserstein distance) and biological metrics (UMAP visualizations, gene ontology annotation). Both works used the DE gene overlap score as the sole evaluation for DE prediction. Mao et al. [22] further showed that model performance is highly context-dependent and drops markedly in cellular context transfer. Others, such as Cole et al. [26], benchmarked foundation-model-derived perturbation embeddings in a KNN regression framework, and found that using interactome- and prior-knowledge-based embeddings often out-perform simple linear baselines in mean root squared error across genetic and chemical perturbations. Beyond model comparisons, several studies have scrutinized evaluation methodology itself. Viñas Torné et al. [25] showed that standard Pearson correlation metrics are dominated by systematic variation — the common, non-perturbation-specific expression shift shared across perturbed cells relative to controls — rather than perturbation-specific signal. Finally, Nicol et al. [35] demonstrated that reusing the same control population to define both predictions and ground truth systematically inflates reported performance.

Across these efforts, evaluation has primarily focused on continuous expression metrics such as Pearson correlation and mean absolute error. The most similar approach to our DE evaluation is the top-*k* DE gene overlap score introduced in Section 1, which measures the overlap between top-*k* predicted and top-*k* true DE genes. This metric, however, does not capture class-specific performance. Our work complements existing studies in two key aspects. First, we focus on DE classification, which accounts for the variance in the expression data and is therefore more aligned with downstream biological analysis. Second, we expand the metric panel beyond DE overlap [13, 23, 24] to include balanced accuracy, macro F1, and class-wise F1, providing a richer characterization of model behavior across the three DE outcome classes. Finally, our experimental splits are derived from the same cell-line datasets as prior work [13, 16, 26], enabling direct comparison of absolute performance while contributing new findings on the classification axis.

## 2 Results

### 2.1 Benchmarking Results for Unseen Perturbation Generalization

Overall, scVI sampling is the best model, and tested non-deep-learning methods are competitive with, and often outperform, the other deep learning models. In this evaluation, scVI sampling achieves the highest macro-F1 on three of the four datasets (K562: 0.588, RPE1: 0.534, HepG2: 0.528, Jurkat: 0.444), placing second to 1-NN (based on GenePT embedding distance) on Jurkat. Across macro-F1, balanced accuracy, and per-class F1, scVI_sampling consistently ranks among the top two models on nearly all datasets (Figure 2), establishing it as a top deep learning method for DE classification in our benchmark for unseen perturbations.

**Fig. 2:**
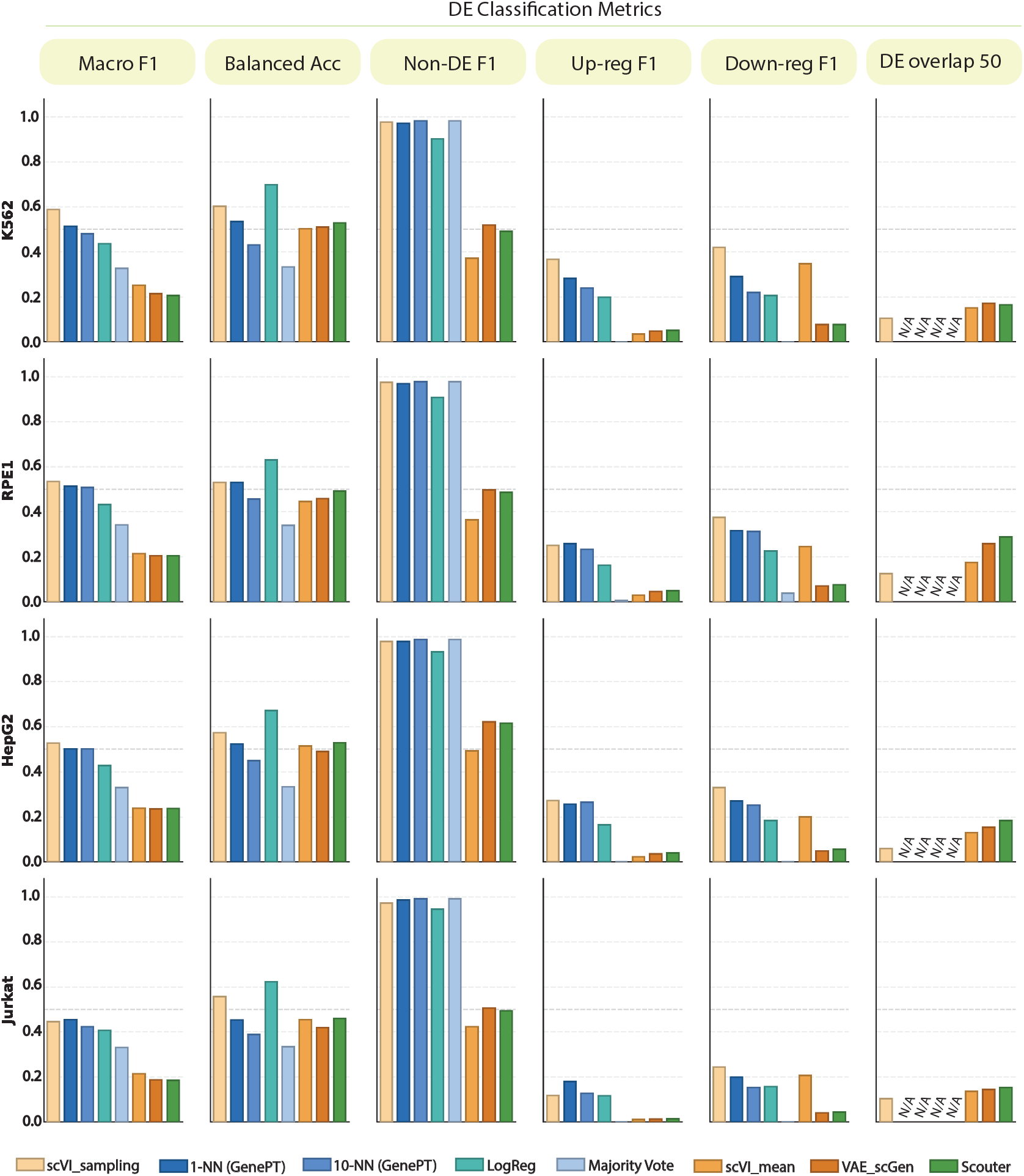
Benchmark results for unseen perturbation generalization. Each subplot shows model scoring on one of the DE classification metrics in a benchmarked dataset. Datasets are represented on the rows, and each column is a DE classification metric. For DE overlap at 50, non-deep-learning models receive ‘NA’ since DE gene ordering requires predicted log_2_ fold change, which classification models don’t have. All subplots received the same model ordering, where models are ranked from highest to lowest based on macro-F1 on DE classification in the K562 dataset. For continuous expression metrics on the unseen perturbation generalization task; see Figure C3.

Among the non-deep-learning methods, 1-NN and regularized logistic regression stand out, matching scVI_sampling on several metrics. 1-NN outperforms scVI_sampling on up-regulation F1 in RPE1 and Jurkat, and otherwise trails the best model by only a small gap (< 13% across all DE classification metrics and datasets). Regularized logistic regression leads on balanced accuracy with an average of 65.6% (K562: 70.0%, RPE1: 63.1%, HepG2: 67.3%, Jurkat: 62.2%), exceeding the second best model by 9% on average.

We further investigate class-wise performance and find that the strong performance of non-deep-learning methods comes from a more aggressive DE-class prediction strategy: logistic regression labels approximately 14.7% of perturbation-gene pairs as DE against a true prevalence of only 2.9% (Table 1), yielding high minority-class recall (average up-regulated: 54.3%; average down-regulated: 55.9%) at the cost of low precision (average up: 9.4%; average down: 11.7%). Sampling-based scVI, by contrast, closely tracks the true DE prevalence (4.3% predicted vs. 2.9% true), trading minority-class recall for substantially higher precision.

**Table 1:** Proportion of perturbation-gene pairs predicted as differentially expressed by each method on the unseen perturbation generalization (within-cell-line) task. True DE prevalence is shown for reference.

|  | K562 | RPE1 | HepG2 | Jurkat | Average |
| --- | --- | --- | --- | --- | --- |
| *Non-deep-learning methods* |  |  |  |  |  |
| Majority vote | 0.00% | 0.10% | 0.00% | 0.00% | 0.03% |
| 10-NN | 0.96% | 1.55% | 0.86% | 0.28% | 0.91% |
| 1-NN | 4.27% | 4.63% | 2.98% | 1.69% | 3.39% |
| Logistic regression | 18.48% | 17.05% | 12.88% | 10.45% | 14.72% |
| *Deep learning methods* |  |  |  |  |  |
| scVI_sampling | 4.09% | 3.92% | 3.75% | 5.37% | 4.28% |
| VAE_scGen | 64.58% | 65.91% | 54.43% | 65.65% | 62.64% |
| Scouter | 67.13% | 67.23% | 55.07% | 67.13% | 64.14% |
| scVI_mean | 76.89% | 76.70% | 66.97% | 72.94% | 73.37% |
| *True DE prevalence* | *3.54%* | *3.94%* | *2.36%* | *1.75%* | *2.90%* |

It turns out that sampling-based scVI is an exception and most deep learning methods benchmarked for this task over-predict DE genes: VAE_scGen (62.6% predicted DE), Scouter (64.1%), and scVI_mean (73.4%) all predict DE signals much higher than the true prevalence (Table 1). Since the non-DE class dominates the evaluation set, over-predicting DE leads to lower accuracy than methods that predict less DE overall, an effect that is especially pronounced for low-fold-change perturbations. Table 2 illustrates this quantitatively: after partitioning all perturbation-gene pairs within each dataset into five equal-sized bins (Q1–Q5, with bin boundaries determined independently per dataset) by true absolute log_2_ fold-change, scVI_mean, VAE_scGen, and Scouter achieve less than 45% accuracy in Q1–Q4, where true DE is rare, while scVI_sampling and non-deep-learning methods maintain above 96% accuracy in the same bins. Crucially, these over-predicting models also fail to surpass scVI_sampling or the leading non-deep-learning methods even in Q5, where true DE pairs are concentrated. This over-prediction pattern therefore offers little-to-none gain in the DE-enriched regime while imposing a heavy cost elsewhere, leading to their weak performance.

**Table 2:** Overall 3-class prediction accuracy by quantile of true |log_2_ FC| for the within-cell-line task, averaged across four datasets (K562, RPE1, HepG2, Jurkat). Quantile bins are computed per dataset and averaged, with approximate ranges shown in column headers. Q1 contains pairs with the smallest absolute expression changes and Q5 the largest.

|  | **Q1**<br>(0–0.016) | **Q2**<br>(0.016–0.034) | **Q3**<br>(0.034–0.060) | **Q4**<br>(0.060–0.108) | **Q5**<br>(>0.108) |
| --- | --- | --- | --- | --- | --- |
| *Non-deep-learning methods* |  |  |  |  |  |
| Majority vote | 99.99% | 99.99% | 99.94% | 99.61% | 85.99% |
| 10-NN | 99.87% | 99.84% | 99.75% | 99.27% | 86.90% |
| 1-NN | 98.76% | 98.62% | 98.22% | 97.02% | 84.65% |
| Logistic regression | 92.36% | 91.55% | 89.60% | 84.96% | 70.58% |
| *Deep learning methods* |  |  |  |  |  |
| scVI_sampling | 98.11% | 97.94% | 97.50% | 96.40% | 86.55% |
| scVI_mean | 14.33% | 16.07% | 20.37% | 30.44% | 55.29% |
| VAE_scGen | 23.36% | 25.79% | 31.60% | 43.48% | 64.41% |
| Scouter | 22.56% | 24.90% | 30.52% | 42.03% | 62.96% |

Lastly, we investigate whether aggressive over-prediction brings an advantage in capturing the most significant DE pairs. We use the metric DE overlap at *k* with *k* = 50, i.e., as the percent overlap between the top-50 true and predicted DE genes. This metric reveals opposite ranking among deep learning methods than other classification metrics: Scouter (avg. 19.7%), VAE_scGen (avg. 18.2%), and scVI_mean (avg. 14.8%) now outperform scVI_sampling (avg. 9.8%) (see Figure 2). This suggests that aggressive over-prediction might benefit models under top-*k* overlap metrics: by assigning DE signals to a larger fraction of genes, models are more likely to rank the most significant true DE genes within their top-50 predictions at the cost of lower precision overall.

We observe the same rank reversal in continuous expression metrics such as mean squared error and Pearson delta, where the best performing model in DE classification has the highest MSE and lowest correlation with the ground truth (Figure C3). Together, these findings demonstrate that no single metric captures the full picture of perturbation prediction performance, and that improved continuous-expression reconstruction, such as lower MSE, does not necessarily produce more accurate DE classifications.

### 2.2 Benchmarking Results for Unseen Cellular Context Generalization

Similar to the unseen perturbation task, on cellular context generalization distinct metrics reveal different model rankings, and a bulk of the deep learning methods do not show better performance in classifying DE status over non-deep-learning counterparts despite their training cost and complexity. 1-NN and STATE are the top performing non-deep-learning and deep learning methods respectively in this task (Figure 3), where 1-NN achieves the highest macro-F1 on K562 and Jurkat (0.477 and 0.469, respectively), and STATE achieves the highest macro-F1 on RPE1 and HepG2 (0.522 and 0.577, respectively). We also evaluate whether the two most accurate methods produce better predictions through calibrated proportions of DE genes after perturbation. We find that the two methods align closely in predicted DE proportions and accuracy by signal strength (Table 3 and Table 4), and their predicted DE proportions are the closest to the true DE prevalence across all four datasets. This reinforces our finding from the unseen perturbation task that calibrated DE proportion is a strong driver of downstream classification performance.

**Table 3:**
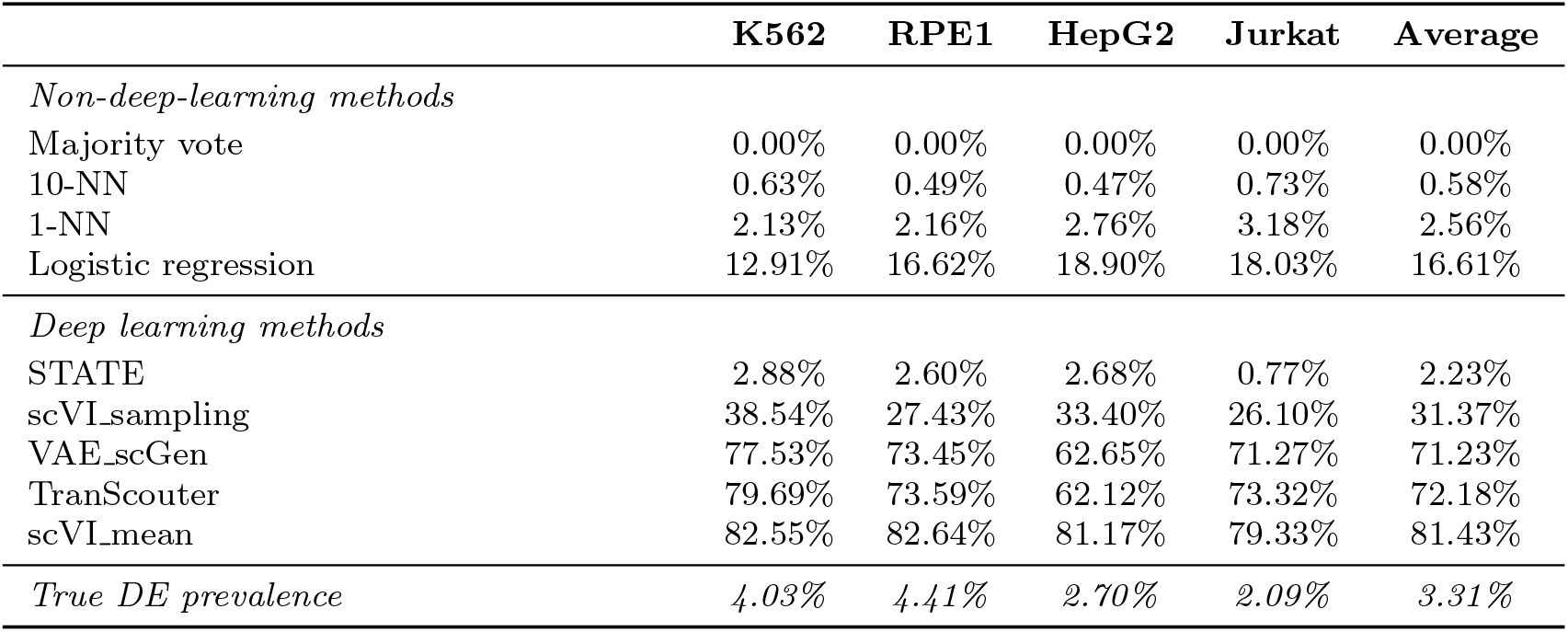
Proportion of perturbation-gene pairs predicted as differentially expressed by each method on the cellular context generalization (cross-cell-line) task.

|  | **K562** | **RPE1** | **HepG2** | **Jurkat** | **Average** |
| --- | --- | --- | --- | --- | --- |
| *Non-deep-learning methods* |  |  |  |  |  |
| Majority vote | 0.00% | 0.00% | 0.00% | 0.00% | 0.00% |
| 10-NN | 0.63% | 0.49% | 0.47% | 0.73% | 0.58% |
| 1-NN | 2.13% | 2.16% | 2.76% | 3.18% | 2.56% |
| Logistic regression | 12.91% | 16.62% | 18.90% | 18.03% | 16.61% |
| *Deep learning methods* |  |  |  |  |  |
| STATE | 2.88% | 2.60% | 2.68% | 0.77% | 2.23% |
| scVI_sampling | 38.54% | 27.43% | 33.40% | 26.10% | 31.37% |
| VAE_scGen | 77.53% | 73.45% | 62.65% | 71.27% | 71.23% |
| TranScouter | 79.69% | 73.59% | 62.12% | 73.32% | 72.18% |
| scVI_mean | 82.55% | 82.64% | 81.17% | 79.33% | 81.43% |
| *True DE prevalence* | *4.03%* | *4.41%* | *2.70%* | *2.09%* | *3.31%* |

**Table 4:** Overall 3-class prediction accuracy by quantile of true |log_2_ FC| for the cross-cell-line task, averaged across four datasets (K562, RPE1, HepG2, Jurkat). Quantile bins are computed per dataset; approximate ranges are shown in column headers.

|  | **Q1**<br>(0–0.017) | **Q2**<br>(0.017–0.037) | **Q3**<br>(0.037–0.065) | **Q4**<br>(0.065–0.116) | **Q5**<br>(>0.116) |
| --- | --- | --- | --- | --- | --- |
| *Non-deep-learning methods* |  |  |  |  |  |
| Majority vote | 100.00% | 99.99% | 99.93% | 99.46% | 84.08% |
| 10-NN | 99.86% | 99.83% | 99.72% | 99.12% | 84.38% |
| 1-NN | 99.07% | 98.94% | 98.60% | 97.51% | 83.67% |
| Logistic regression | 89.03% | 88.30% | 86.50% | 82.65% | 68.43% |
| *Deep learning methods* |  |  |  |  |  |
| STATE | 99.36% | 99.26% | 98.99% | 97.97% | 85.81% |
| scVI_sampling | 67.67% | 68.48% | 69.60% | 70.38% | 67.55% |
| VAE_scGen | 19.70% | 21.48% | 25.53% | 33.26% | 48.96% |
| TranScouter | 19.71% | 21.38% | 25.20% | 32.28% | 46.21% |
| scVI_mean | 11.00% | 12.12% | 14.83% | 20.62% | 38.13% |

**Fig. 3:**
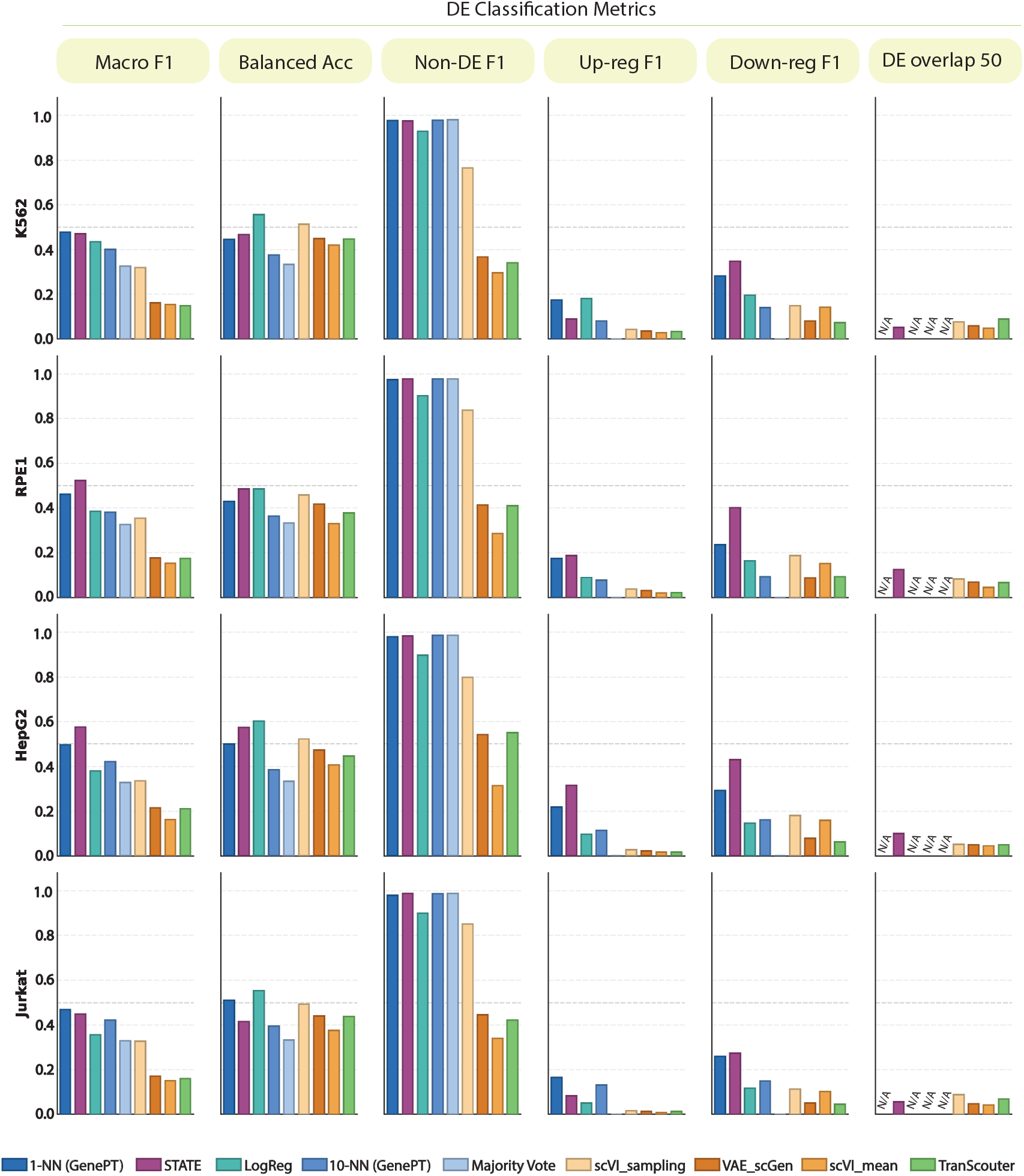
Benchmark results for cellular context generalization. Each subplot shows model scoring on one of the DE classification metrics in a benchmarked dataset. Datasets are represented on the rows and each column is a DE classification metric. For DE overlap at 50, non-deep-learning models receive ‘NA’ since DE gene ordering requires predicted log_2_ fold change, which classification models don’t have. All subplots received the same model ordering, where models are ranked from highest to lowest based on macro-F1 on DE classification in the K562 dataset. For continuous expression metrics on the cellular context generalization task see Figure D4.

Notably, scVI_sampling, the best performer in the within-cell-line task, substantially over-predictes DE pairs in cross-cell-line transfer (26–39% vs. a true prevalence of 2–4%), suggesting that different generalization tasks may favor different model characteristics. Both 10-NN and regularized logistic regression outperform TranScouter, VAE_scGen, and scVI_mean on macro-F1 and reveal predicted DE proportions closer to the true DE prevalence (Figure 3, Table 3), echoing with the finding in within-cell-line task that non-deep-learning methods remain competitive in DE predictions compared to deep learning models.

Unlike within-cell-line prediction, the best performing deep learning model in DE classification, STATE, also shows leading performance in the majority of continuous expression metrics (Figure D4). However, we note that STATE is the worst-performer in per-gene Pearson correlation, which is defined as the Pearson correlation between a gene’s predicted and observed mean expression profile across all perturbations, averaged over genes (see Methods at section 3 for details). This could suggest that STATE fails to reproduce the gene-specific perturbation-response rankings that per-gene Pearson correlation measures. Overall, this observation adds to our previous finding that a diverse metric panel is needed to fully assess perturbation models.

### 2.3 Benchmarking Results for Post-Prediction Sparsity Calibration

Motivated by the observation that most single-cell perturbation models produce predicted expression matrices denser than the ground truth, we apply a sparsity calibration post-prediction to align predicted expression sparsity to that of the unperturbed control cells (Figure 4a). In the within-cell-line setting, three of the four deep learning models (Scouter, VAE_scGen, and scVI_mean) show gains in multiple metrics including balanced accuracy, macro-F1, and per-class F1 post-calibration (Figure E5). Macro-F1 improvements range from +0.065 (relative change: +32%) to +0.237 (relative change: +99%), with the biggest improvement being scVI_mean on prediction of unseen perturbations in HepG2 (Figure 4b). The same pattern holds for cross-cell-line evaluation, where TranScouter, VAE_scGen, and scVI_mean again respond positively (macro-F1 improvements from +0.037 to +0.172) (Figure 4c).

**Fig. 4:**
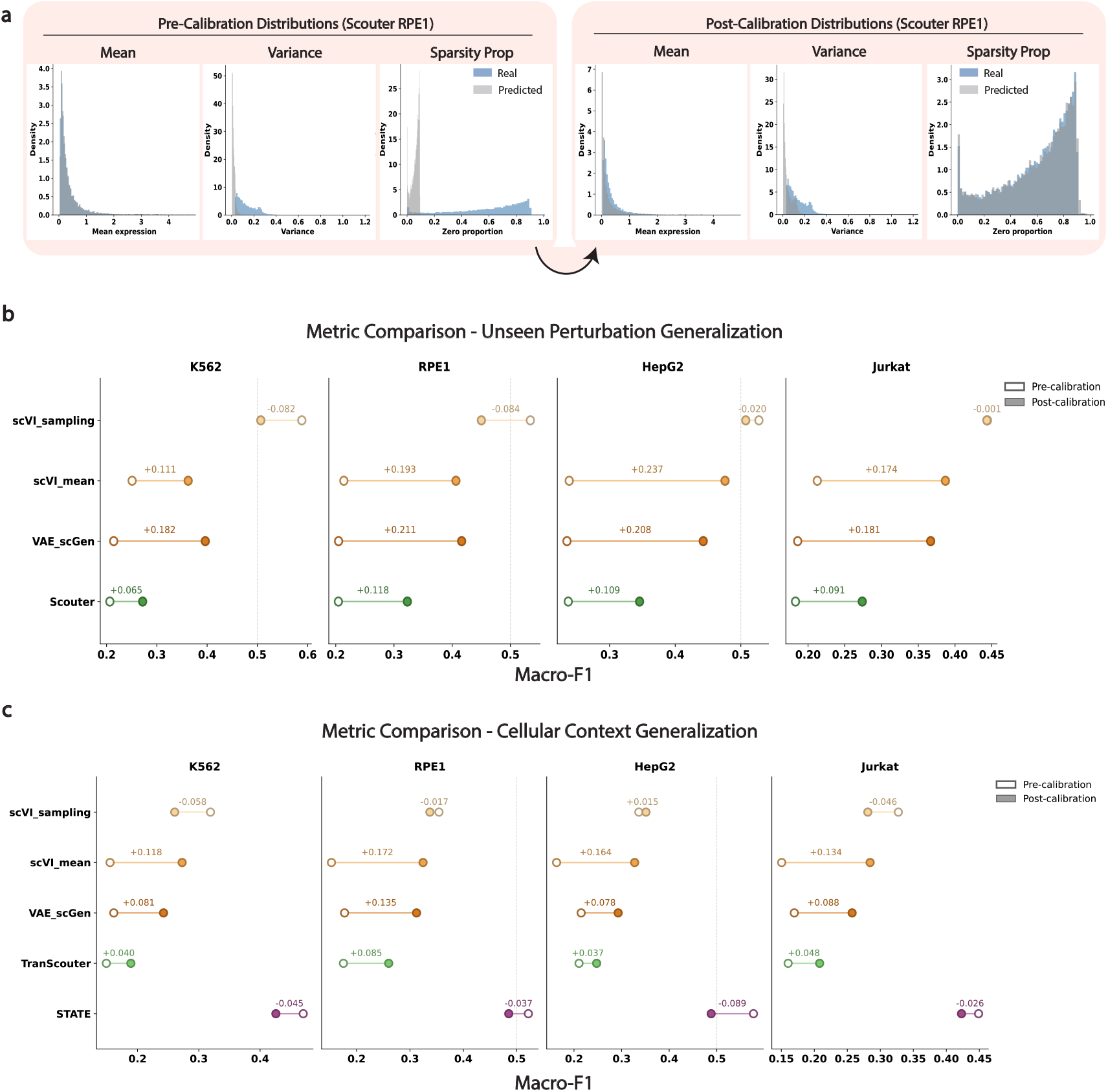
Pre- and Post-sparsity calibration effect for unseen perturbation generalization (within-cell-line prediction) and cellular context generalization (cross-cell-line prediction). **a)** Pre-calibration per-gene expression mean, variance, and sparsity level for Scouter on the RPE1 dataset. Grey distributions are plotted based on Scouter’s predictions and blue distributions are from the ground truth dataset. These are compared with post-sparsity calibration distributions to show distributional alignments. **b)** Change in macro-F1 of the within-cell-line prediction task post-sparsity calibration for models in all benchmarked datasets. **c)** Change in macro-F1 of the cross-cell-line prediction task post-sparsity calibration.

By contrast, scVI_sampling and STATE show negligible or slightly negative changes in both evaluation scenarios. We observe that prior to calibration, their predicted expression distributions are already closely aligned to that of the control cells, which likely arise from their individual model design: STATE uses Maximum Mean Discrepancy (MMD) loss [36] during training that matches the empirical distribution of predicted single-cell expression vectors to that of the observed cells, and scVI_sampling explicitly learns a Zero-Inflated Negative Binomial (ZINB) distribution with a dedicated zero-inflation parameter [11]. We therefore hypothesize that sparsity calibration is a useful post-prediction technique as well as a notable factor in model design. Specifically, when the underlying model does not account for the sparsity in the single-cell data, calibration could boost a perturbation model’s DE prediction performance.

With closer inspection, we find that across all improved models, the gain of calibration is most pronounced in non-DE F1 (see Figure E5). In other words, calibration primarily corrects for over-prediction of DE events, shifting predicted class proportions closer to the true distribution in which non-DE genes overwhelmingly dominate, consistent with what we have observed in previous benchmarking tasks.

### 2.4 Benchmarking Results of Models on Curated Subsets

Motivated by the growing use of curated perturbation datasets to account for class imbalance (i.e., most genes remain non-DE post-perturbation) [16–18], we compare model performance on the full dataset and on subsets curated by Wu et al. [16], which downsamples non-DE pairs. We find that nearly all models show substantial improvements in DE classification performance. Across both within-cell-line and cross-cell-line evaluations, the most pronounced improvements are in non-deep-learning methods, suggesting that classification-based approaches are more sensitive to data curation (Figure 5a, Figure 5b), which is likely because they learn directly from DE labels.

**Fig. 5:**
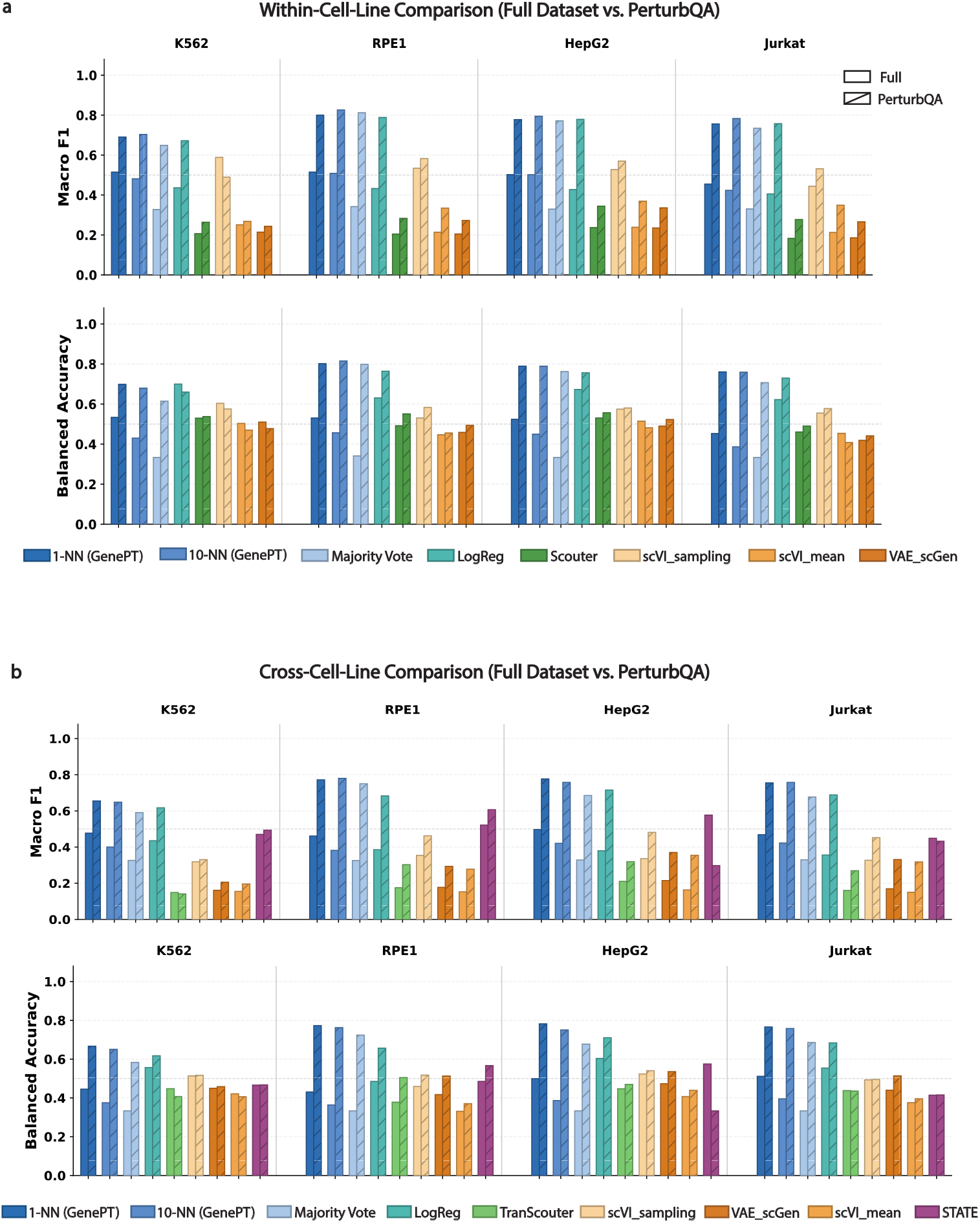
Benchmark results comparing model performance when trained on the full, unsampled dataset vs. down-sampled perturbQA dataset. **a)** Performance comparison on the unseen perturbation generalization task (within-cell-line evaluation). **b)** Performance comparison on the cellular context generalization task (cross-cell-line evaluation).

Deep learning models also show overall improvements in macro-F1 and balanced accuracy, though the magnitude varies by dataset. In the within-cell-line setting, almost all deep learning methods improve in macro-F1 across four datasets, with the sole exception of scVI_sampling in K562 (Figure 5a). In the cross-cell-line setting, macro-F1 is broadly higher on the curated subset, with exception of STATE on HepG2 and Jurkat (Figure 5b). Compared with non-deep-learning approaches, these gains are generally more modest, suggesting that deep learning methods, which learn from continuous expression profiles, are less dependent on class balance in the training data. These results underscore a critical consideration for model evaluation: performance metrics are meaningful relative to the data conditions under which a model was trained and evaluated on. Improvements observed on curated subsets should be carefully interpreted as they could be reflecting the consequence of prior data sampling rather than reflecting true model gains.

## 3 Methods

### 3.1 Problem Setup and Evaluation Tasks

Following Phillips et al. [17], we formulate differential expression prediction as a three-class classification problem. Let *g* denotes a gene, *p* a perturbation, and *P*_train_ the set of all training perturbations. Let *y*_*g,p*_ be the observed DE status of gene *g* under perturbation *p*, and *ŷ*_*g,p*_ the predicted DE class. The task is to predict whether *ŷ*_*g,p*_ is: (1) non-differentially expressed (Non-DE), (2) up-regulated (Up), or (3) down-regulated (Down). We evaluate on two downstream tasks:

#### Unseen Perturbation Generalization

This task evaluates how well a model can predict DE outcomes under an unseen perturbation from the same cell line of training samples (i.e., within-cell-line evaluation). In this task, the dataset from a cell line is split by perturbations into a 60% training and a 40% testing set, so that all perturbations in the test set are unseen during training. Dataset statistics are provided in Table 5. For all benchmarked methods, we train on this same train split and report performance on the same 40% test set (see training schematic in Figure 1) to ensure fair model comparisons.

**Table 5:** Per-dataset DE statistics. “M” indicates the unit of the measurement is in millions.

|  | **K562** | **RPE1** | **HepG2** | **Jurkat** |
| --- | --- | --- | --- | --- |
| Total pairs (M) | 17.61 | 20.93 | 23.03 | 21.25 |
| Non-DE (%) | 96.61 | 95.96 | 97.61 | 98.35 |
| Up-reg (%) | 1.44 | 1.51 | 0.99 | 0.50 |
| Down-reg (%) | 1.95 | 2.53 | 1.40 | 1.15 |
| Train pairs (M) | 10.56 | 12.55 | 13.81 | 12.74 |
| Test pairs (M) | 7.05 | 8.38 | 9.22 | 8.51 |
| Perturbations | 2,057 | 2,393 | 2,393 | 2,393 |
| Genes | 8,561 | 8,748 | 9,623 | 8,881 |

#### Cellular Context Generalization

This task evaluates how well a model can predict DE outcomes when provided a perturbation it has seen during training but in another cell line (i.e., cross-cell-line evaluation). Given a held-out cell line, all models are trained using perturbation-gene pairs from the other training cell lines and we report performance on the 40% test set of the held-out cell line (see training schematic in Figure 1). We train on the common gene space across four cell lines (6, 641 genes) to avoid introducing 0 expression for unobserved perturbation-gene pairs into the training data. For STATE [13] specifically, we include the 60% training set of the held-out cell line during model training (see training schematic in Figure 1) following training convention in the official codebase [37], so it is worth noting that STATE is performing a “few-shot” prediction rather than a “zero-shot” prediction in the sense that the referred version requires some observations in the new cell line.

### 3.2 Benchmarked Datasets and Processing

#### Dataset Processing

We benchmark on essential gene screens from K562 [32], RPE1 [32], Jurkat [33], and HepG2 [33] cell lines, chosen for their widespread use in perturbation modeling. For each dataset, we followed prior work [8, 13, 16] and normalized raw UMI counts by dividing by the total counts per cell, scaling to 10,000, and applying a log(1 + *x*) transformation.

#### DE Statistics

DE statistics are obtained through Wilcoxon rank-sum tests followed by Benjamini-Hochberg correction per perturbation. A pair (*p, g*) is considered DE if the adjusted p-value is less than 0.01. The sign of the log-fold change further determines up- or down-regulation. We employ this same procedure to obtain ground truth DE labels from observed expressions, and predicted DE labels from expressions predicted by deep learning models.

#### Full vs. Curated Dataset

For each benchmarked cell line we train and evaluate models on a full and a curated subset. First, we evaluate models on **PerturbQA-downsampled datasets** where in Wu et al. [16] non-DE pairs were downsampled to address the severe class imbalance present across all four datasets (95-98% non-DE, see Table 5), which can bias both training and evaluation, and DE pairs were restricted to those with top statistical significance. Second, we evaluate on the **full dataset without downsampling**, reflecting the more realistic setting where no prior knowledge of class distribution is available. We annotate our results to distinguish between these two settings. In evaluating the downsampled dataset, we used the train-test split defined by Wu et al. [16].

### 3.3 Benchmarked Methods

#### 3.3.1 Perturbation Majority Vote

We first consider a simple majority vote rule based on the hypothesis that a gene’s response to perturbation largely reflects its intrinsic properties rather than the specific perturbation applied. That is, if gene *g* tends to be up-regulated across most training perturbations, we predict it will also be up-regulated under a new, unseen perturbation. Formally, for each gene *g*, we assign the most frequent response class observed across all training perturbations where 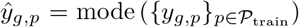, *y*_*g,p*_ ∈ {Up, Down, Non-DE}. Because this baseline ignores perturbation identity entirely, it produces the same prediction for gene *g* regardless of which perturbation is applied. Models that fail to outperform this baseline may not be capturing meaningful perturbation-specific signal.

#### 3.3.2 Weighted *K*-Nearest Neighbors

To incorporate perturbation-specific information, we use a weighted *K*-nearest neighbors (KNN) classifier with GenePT embeddings [30] to represent perturbations. For each query pair (*p, g*), we identify the *K* most similar training perturbations via cosine similarity in embedding space and predict *g*’s response through distance-weighted voting. This method evaluates whether local neighborhoods in perturbation embedding space capture sufficient information for predicting gene responses. Our primary results used *K* ∈ {1, 10}; a larger value of *K* typically leads to suboptimal performance. We also conduct a sensitivity analysis comparing KNN classifiers trained using alternative embeddings and find consistent performance across embedding choices (see Appendix A).

#### 3.3.3 Regularized Logistic Regression

We then ask whether a parametric linear classifier trained on the same embeddings can learn broader, transferable patterns of gene response. For each gene *g*, we train an *ℓ*_2_-regularized multinomial logistic regression model that takes the GenePT embedding [30] of a query perturbation as input and predicts the DE class (with regularization strength = 0.01). Let **y**_*g*_ has each entry *y*_*g,p*_ denotes the observed DE class of gene *g* under perturbation *p*, **W** the matrix of embedding vectors where each row represents a perturbation, the logistic model finds parameters ***θ*** such that logit(**y**_*g*_ ) = **W*θ***. The formulation is adapted from GenePert [10] and directly tests whether linear decision boundaries in embedding space generalize beyond what local neighborhood retrieval affords.

#### 3.3.4 Variational Autoencoder

We adapt scGen [12] to predict perturbation effects for both within-cell-line and cross-cell-line settings (see Figure B2 for adapted model architecture). Denoting **z**_*p*_ the low-dimensional latent representation of a cell under perturbation *p*, during training, the model reconstructs training cell expressions while conditioning on perturbation identity through the GenePT embeddings. For each perturbation *p* ∈ *P*_train_, the model estimates its latent perturbation effect ***δ***_*p*_ as the difference between the average latent representations of the perturbed cells and control cells (i.e., ***δ***_*p*_ = avg(**z**_*p*_) − avg(**z**_ctrl_)). In the within-cell-line setting, to predict an unseen perturbation *p*, we select the *k* most similar training perturbations based on cosine similarity of GenePT embeddings:

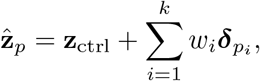

where *w*_*i*_ is proportional to the cosine similarity between *p* and the *i*-th selected perturbation. In the cross-cell-line setting, each 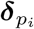 is first averaged across training cell lines before the weighted combination:

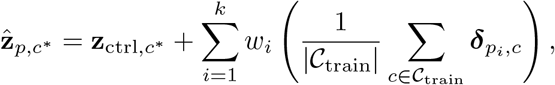

where *c*^∗^ denotes the held-out target cell line, *C*_train_ denotes the set of training cell lines, and 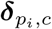 represents latent perturbation effect of perturbation *p*_*i*_ estimated from cell line *c*. In both settings, the predicted gene expression profile is obtained by decoding 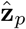 through the decoder.

#### 3.3.5 scVI (Mean and Sampling)

We further benchmark on scVI [11] which is an autoencoder-based method that models gene expression distributions taking into account technical noise and batch effects. We adapt STATE’s benchmarking implementation of scVI [13, 38], which takes a control-cell expression profile and a perturbation identity as input, and outputs the predicted perturbed gene expression profile. Basal expression profiles from control cells are encoded into a low-dimensional latent representation using a variational encoder. Perturbation identities are represented with GenePT embeddings and concatenated with the latent basal state, together with available technical covariates such as batch labels. A decoder then maps this combined representation to the parameters of a zero-inflated negative binomial distribution over genes, allowing the model to capture both overdispersion and excess zeros in single-cell count data. We report two forms of model output: one based directly on the mean of the learned output distribution (scVI_mean), and the other based on expression profiles sampled from that distribution (scVI_sampling).

#### 3.3.6 Scouter and TranScouter

To include a more diverse set of models with varying loss functions we benchmark on Scouter [14] and TranScouter [31], both use a compressor-generator architecture and an autofocus direction-aware loss initially proposed in GEARS [8]. Scouter is designed for within-cell-line prediction, generating transcriptional responses to genetic perturbations within the same cell line used for training. TranScouter extends Scouter to the cross-cell-line setting through two modifications: an encoded cell line representation is reintroduced near the decoder bottleneck to preserve cell-line-specific information during decoding, and condition-aware pairing is applied during training so that each perturbed cell is matched only with control cells from the same cell line, rather than pooled freely across conditions. Given a perturbation, both models take as input the log-normalized gene expression profile of a randomly sampled control cell and the LLM-based embedding of the perturbed gene, and output the predicted expression profile of a perturbed cell.

#### 3.3.7 STATE

STATE [13] is a transformer-based model that predicts perturbation-induced gene expression shifts at the population level by operating on sets of cells rather than individual observations. The core State Transition (ST) model constructs a composite additive representation **H** = **H**_cell_ + **H**_pert_ + **H**_batch_ for each cell set, where **H**_cell_ encodes control expression profiles, **H**_pert_ encodes perturbation identity via a learned embedding, and **H**_batch_ encodes batch covariates. The ST model produces output **O** = **H** + *f*_ST_(**H**) via a residual connection over a transformer backbone, and is trained to minimize a MMD loss between predicted and observed distributions. We use STATE in the gene expression mode (without the pretrained State Embedding component), operating directly on the common gene universe across all four cell lines; see Appendix B for details on training procedures, including hyperparameters.

### 3.4 Sparsity Calibration

We apply a post-hoc, gene-level zero calibration to align the sparsity of predicted expression with that observed in unperturbed control cells. For each gene *g*, we compute the zero-expression fraction *z*_*g*_ across all control cells as an estimate of natural sparsity in the given dataset:

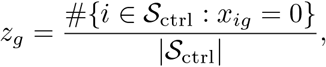

where *S*_ctrl_ denotes the set of control cells, *x*_*ig*_ the expression of gene *g* in cell *i*. A gene-specific calibration threshold *X*_*g*_ is then defined as the *q*_*g*_-th quantile of predicted expression values pooled across all perturbed cells, where *q*_*g*_ = *r* · *z*_*g*_ and *r* ∈ [0, 1] is a global scaling factor. Predicted expression values at or below *X*_*g*_ are set to zero; all other values and control cell measurements are left unchanged. We set the scaling factor to *r* = 1, which postulates that the sparsity level before and after perturbation should remain the same.

### 3.5 Evaluation Metrics

#### 3.5.1 DE Classification Metrics

##### Per-Class F1

For each DE class *j* ∈ {Up, Down, Non-DE}, precision and recall are defined as

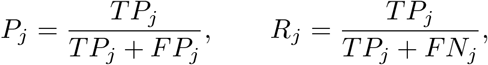

where *TP*_*j*_, *FP*_*j*_, and *FN*_*j*_ denote true positives, false positives, and false negatives for class *j*, respectively. The per-class F1 score is calculated as

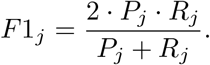

##### Macro F1

Macro F1 averages the per-class F1 scores equally across all *J* = 3 classes, giving equal weight to each class regardless of its frequency:

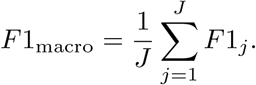

##### Balanced Accuracy

Balanced accuracy averages per-class recall across all classes, correcting for class imbalance:

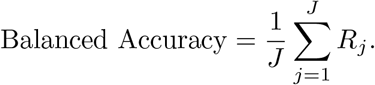

#### 3.5.2 Continuous Expression Metrics

The following metrics operate on mean gene expression profiles (pseudobulks) computed per perturbation. Let *T* denote the number of perturbations, *G* the number of genes, 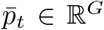 the predicted pseudobulk for perturbation *t, p*_*t*_ ∈ ℝ^*G*^ the observed pseudobulk, and *c* ∈ ℝ^*G*^ the control pseudobulk. DE overlap at 50, mean absolute error (MAE), mean squared error (MSE), and Pearson delta are all metrics originally proposed in Adduri et al. [13].

##### DE Overlap at 50

For each perturbation *t*, the top-*k* differentially expressed genes are selected from the true and predicted DE gene sets respectively by ranking all DE genes in descending order of absolute log_2_ fold change. The overlap score at *k* = 50 is

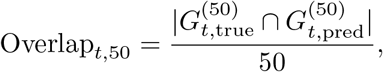

where 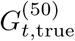 and 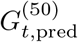 denote the respective top-50 DE gene sets. The final metric is the mean over all perturbations with at least one true DE gene.

##### Mean Absolute Error

MAE measures the average per-perturbation magnitude of expression prediction errors across all genes:

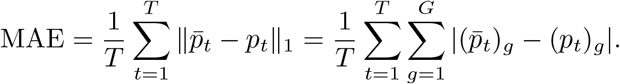

##### Mean Squared Error

MSE penalizes large per-gene prediction errors quadratically:

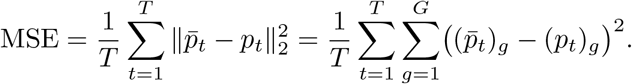

##### Pearson Global Correlation

A single Pearson correlation coefficient is computed over all (*T* × *G*) perturbation-gene pairs by flattening both the predicted and observed pseudobulk matrices:

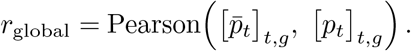

This provides a global measure of linear agreement between predicted and observed expression levels across the entire perturbation-gene space.

##### Pearson Delta Correlation

Following Adduri et al. [13], we compute expression deltas as the element-wise absolute deviation of each perturbation pseudobulk from the control: 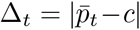 (predicted) and 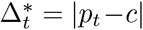 (observed), where |·|denotes the element-wise absolute value. The metric is the Pearson correlation computed jointly over all (*T* × *G*) entries:

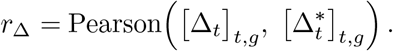

This captures how accurately a model reproduces the magnitude of perturbation-induced expression shifts relative to the unperturbed baseline.

##### Pearson Per-Gene Correlation

For each gene *g*, a Pearson correlation coefficient is computed between the predicted and observed mean expression of that gene across all *T* perturbations:

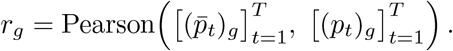

Genes for which either the predicted or observed perturbation response vector is constant are excluded. The final metric is the mean 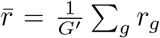 over the *G*′ retained genes. This evaluates whether the model correctly captures which perturbations up- or down-regulate each individual gene.

## 4 Discussion

In this study, we presented a systematic benchmark of five deep learning expression models and three non-deep-learning classification methods on differential expression classification across four human cell-line datasets and two generalization regimes. Our results demonstrate that simple classification methods, in particular nearest-neighbor approaches with perturbations represented by LLM-derived embeddings, are broadly competitive with and often outperform deep learning models on DE classification. Moreover, no single method uniformly outperforms the rest across all datasets and metrics. We further showed that better performance on continuous expression metrics does not equate better DE classification performance, highlighting the need to evaluate DE prediction directly rather than relying on continuous reconstruction metrics alone. We also found that post-prediction sparsity calibration emerged as an effective and practical correction for deep learning models that do not internally account for expression sparsity. A simple calibration based on control cell sparsity level consistently improved DE classification performance without requiring retraining. Finally, evaluation on a curated subset with downsampled non-DE pairs revealed that dataset composition profoundly shapes reported performance, particularly for classification-based baselines, reinforcing the need to interpret benchmark results in the context of the data distribution on which models are trained and assessed.

Our current work is subject to a few limitations. First, we restrict our benchmark to four commonly used immortalized cancer cell lines in perturbation modeling. These datasets may not be representative of the broader landscape of cellular contexts relevant to biomedicine, including primary cells [39], patient-derived samples [40], and in vivo systems [2, 28]. As more perturbation datasets become available, future work should investigate whether our conclusions generalize to more biologically relevant settings. Second, we evaluate exclusively single-gene perturbations, leaving combinatorial perturbations [8, 34] and chemical perturbations [41] outside the scope of this benchmark. A similar benchmark study on chemical perturbations could provide insight into the comparison of different model classes in that setting and verify whether our findings transfer. Third, the three-class DE label schema we adopt (up-regulated, down-regulated, unchanged) is inherently dependent on the statistical thresholds used to assign labels. Therefore, our findings could change under alternative DE-calling procedures. However, by stratifying our results based on the magnitude of perturbation effects, we partially mitigate this sensitivity concern.

Other directions for future work include benchmarking on perturbation screens from other species [42], and experimenting with recent developments in generative model calibration [43] to assess whether novel calibration procedures that account for the entire distribution of gene expression, instead of just sparsity or mean expression, could provide better results — both for post-hoc calibration and for model training and loss function design. Finally, as the field moves toward evaluating whether models recover coordinated groups of DE genes rather than only classifying each gene independently, it will be important to develop benchmarks that test whether predicted DE patterns identify the same biological pathways, cell-state changes, and gene groups affected by a perturbation. Such evaluations would better align model assessment with the biological questions that motivate perturbation-response prediction.

## Code and data availability

Code used to generate the analysis in this paper can be found at https://github.com/ivysun14/Project-Perturb.

## Appendix A KNN Sensitivity Analysis

To ensure model performance is not driven by embedding choice alone, we evaluate four types of perturbation representations for the KNN classifier. (1) GenePT [30] embeddings are 3,072-dimensional vectors produced by encoding NCBI/UniProt gene descriptions with the OpenAI text-embedding-3-large model [44] and provide full coverage of all perturbations in our datasets. (2) ESM2 embeddings are derived from the ESM2 protein language model applied to canonical protein sequences [45]. (3) WikiCrow embeddings are produced by encoding WikiCrow-generated gene summaries [46] with the same OpenAI text-embedding-3-large model used for GenePT. (4) Co-pathway (co-path) representations are generated using the `gseapy` library [47] in Python. Perturbation similarity is quantified by the Jaccard similarity of shared pathway membership across MSigDB [48], KEGG [49], and Reactome gene sets [50], pre-computed and stored as a ranked neighbor list per perturbation.

These four representation types have different coverage of perturbations in the benchmarked dataset. For coverage details see Figure A1a. To ensure that performance differences across embedding types reflect representation quality rather than differences in the evaluated perturbation set, all sensitivity analysis applies a coverage filter that restricts the evaluated perturbations to the intersection of perturbations covered by all four representation types. Results of the sensitivity analysis can be found in Figure A1b.

**Fig. A1:**
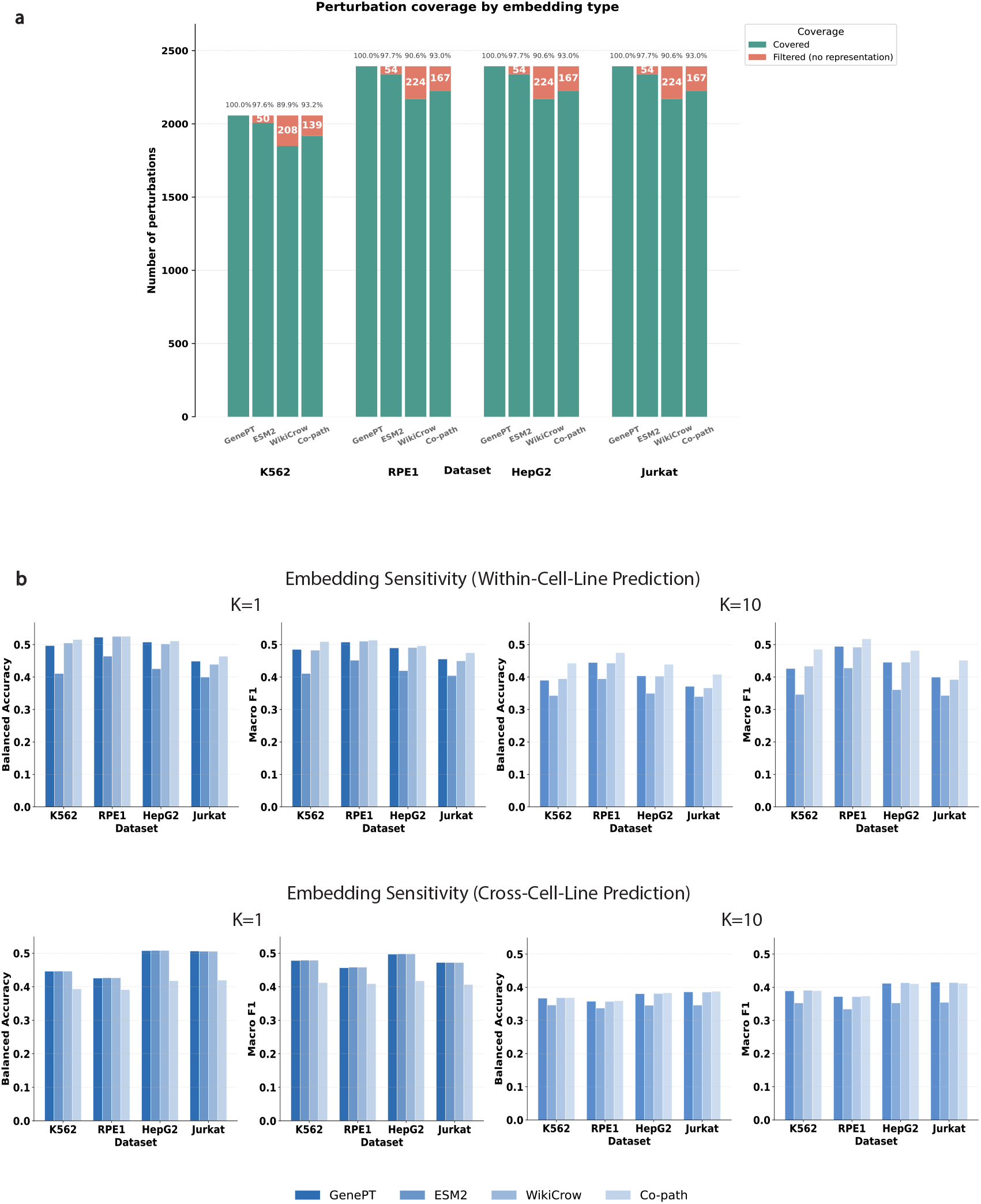
**a)** Perturbation coverage by embedding types. Percentage of perturbation missing representations in each dataset by each type is quantified. **b)** Sensitivity analysis of different embeddings for KNN classifiers in within-cell-line and cross-cell-line evaluation tasks for K=1 and K=10.

## Appendix B Training Specifications and Model Details

### scVI (Mean and Sampling)

These models are trained by minimizing the ZINB negative log-likelihood of the observed perturbed expression profile, together with a KL divergence term that regularizes the latent basal representation toward a standard normal prior. Models are optimized with Adam using mini-batches of 512 cells, a learning rate of 5 × 10^−4^, weight decay of 4 × 10^−7^, gradient clipping at 10. Training runs for up to 25,000 optimization steps, with validation performed every 500 steps.

### Scouter and TranScouter

For each perturbation, the number of predictions generated was matched to the observed cell count for that perturbation in the original dataset. Both models were trained using the Adam optimizer with exponential learning rate decay (factor 0.9 per epoch), early stopping (patience 5, minimum improvement 0.001), and gradient clipping (max norm 1.0). Scouter was trained with a learning rate of 0.001, a batch size of 256, and a maximum of 40 epochs. TranScouter was trained with a learning rate of 0.0001, a batch size of 512, and a maximum of 50 epochs.

### STATE

We trained STATE via the official CLI with cell set size = 64, hidden dimension = 672, batch size = 32, learning rate = 10^−3^, and 50,000 training steps, with batch correction enabled using the sequencing run (`gem_group`) as the batch covariate. A random 10% of the target cell line’s training perturbations was reserved as a validation set. Perturbations absent from the training perturbation universe were identified and excluded from evaluation since STATE is meant for context generalization and cannot predict on a completely unseen perturbation.

**Fig. B2:**
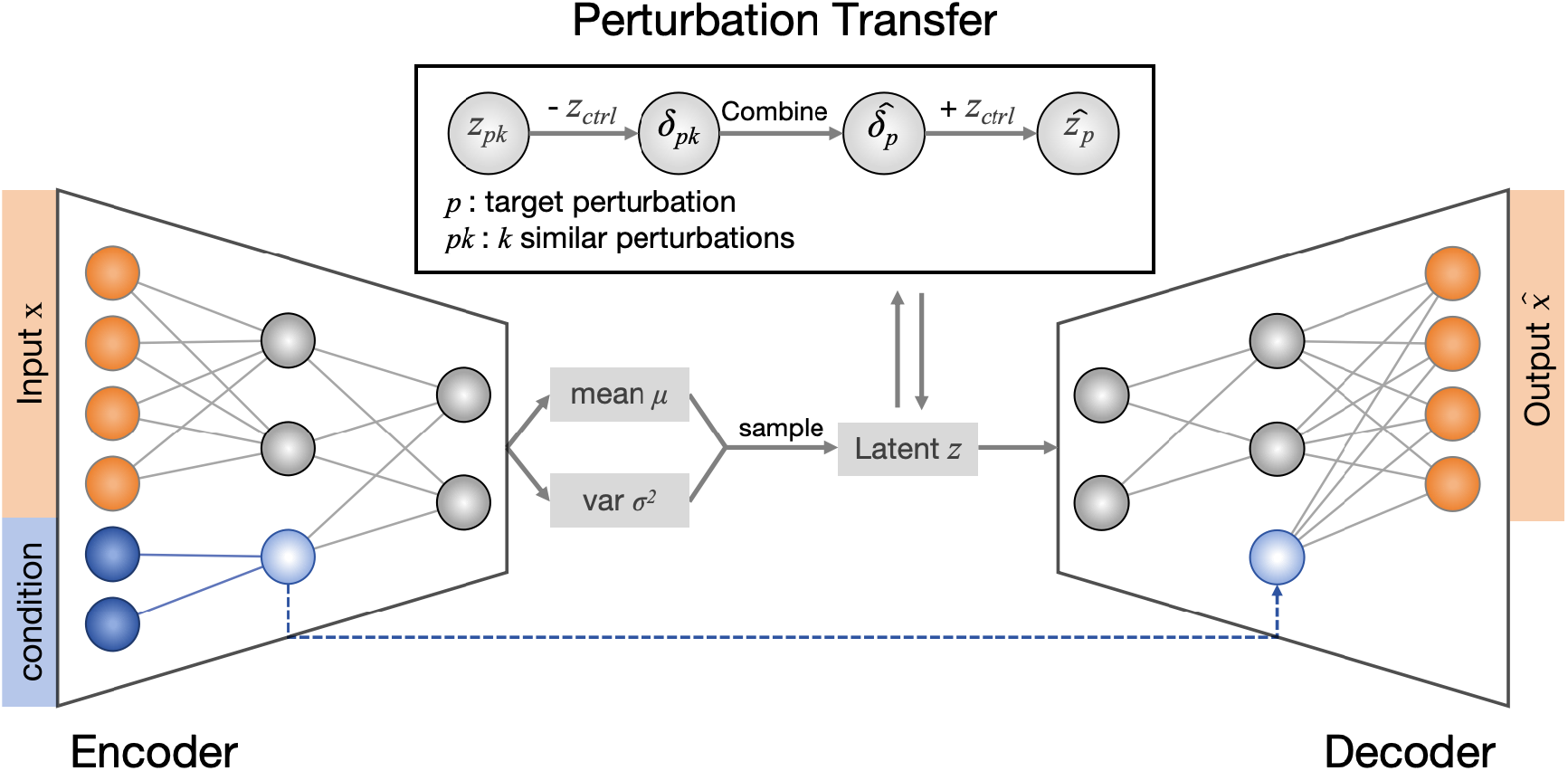
Overview of the adapted scGen model architecture. Perturbation conditions represented by embedding vectors are concatenated to the input expression profile and passed to the encoder. The decoder aims to recover input expression profile, through which the model learns a delta specific to the perturbation that captures expression profile change in the latent space.

## Appendix C Model Ranks on Continuous Expression Metrics — Unseen Perturbation Generalization

**Fig. C3:**
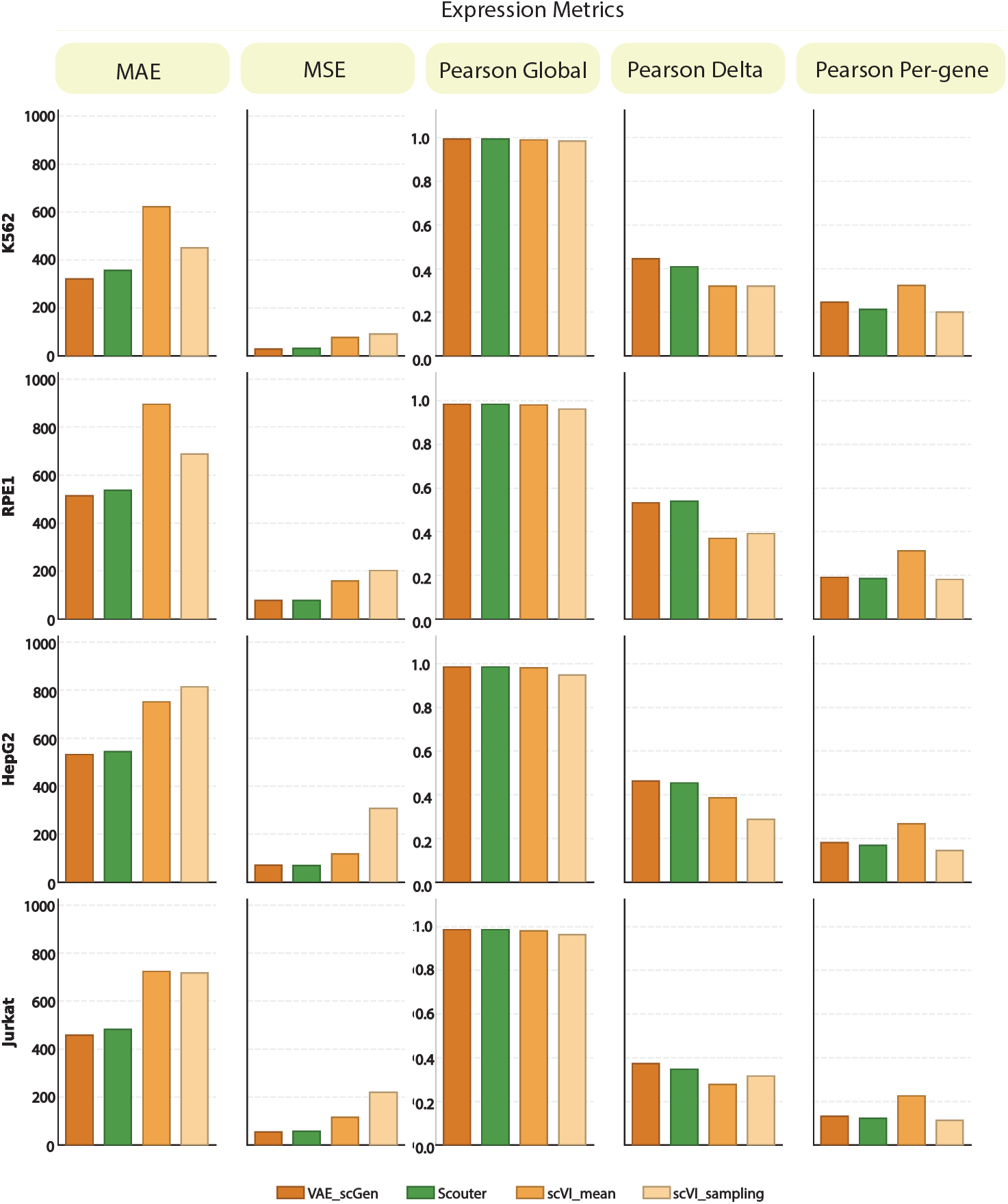
Expression-based metrics for unseen perturbation generalization task.

## Appendix D Model Ranks on Continuous Expression Metrics — Cellular Context Generalization

**Fig. D4:**
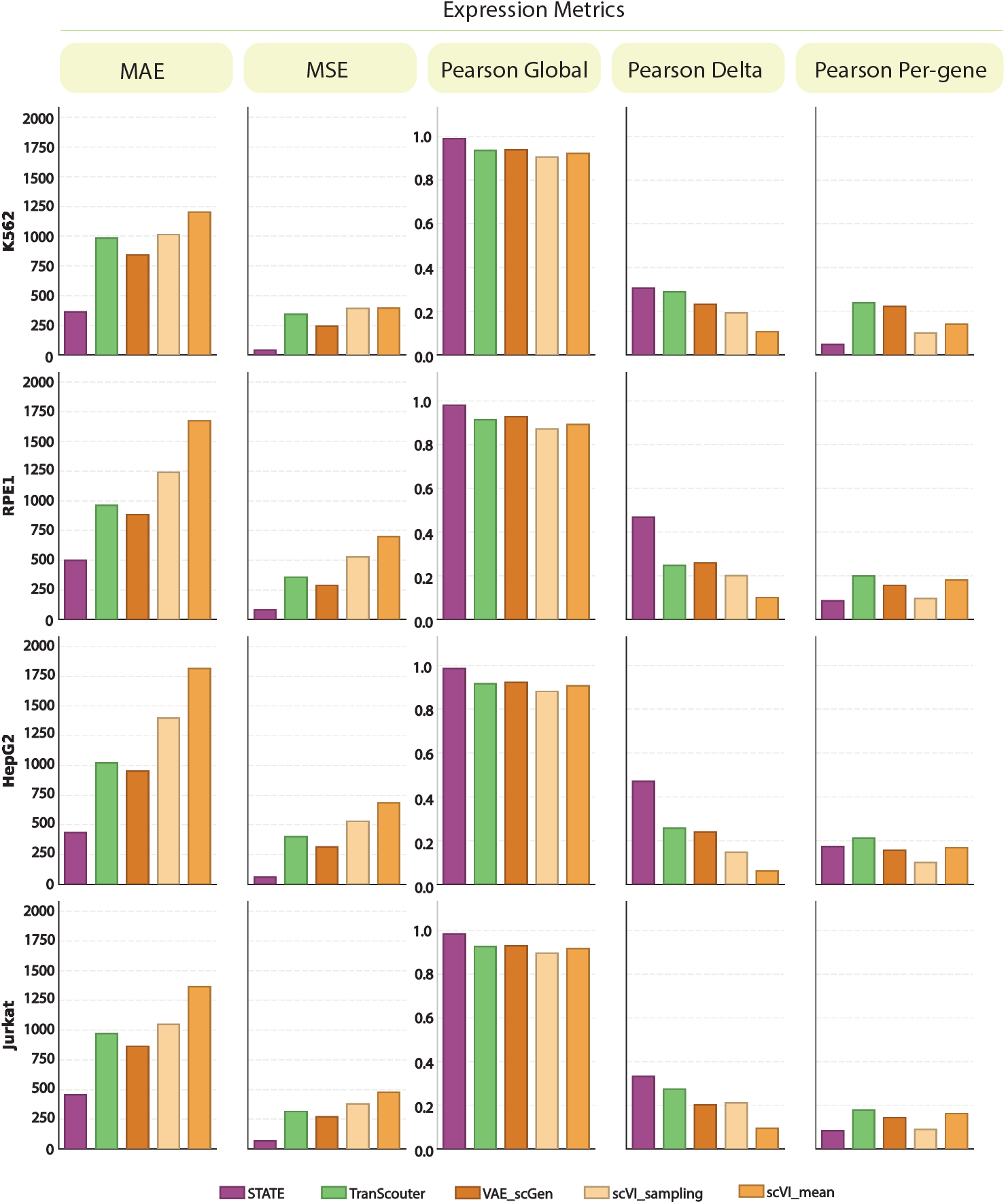
Expression-based metrics for cellular context generalization task.

## Appendix E Post-Sparsity-Calibration Metric Change

**Fig. E5:**
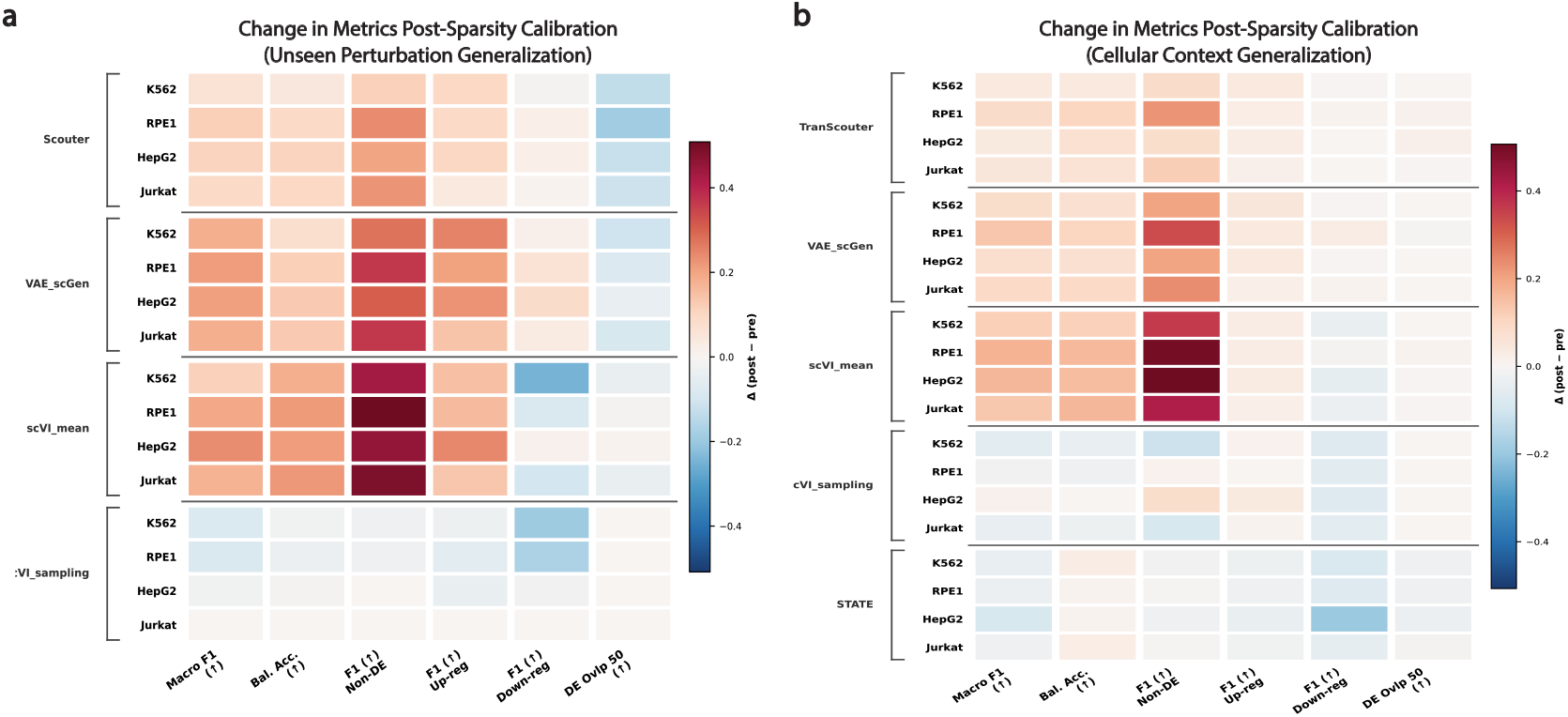
**a)** Change in DE classification metrics post-sparsity calibration for unseen perturbation generalization. Calibration delta (Δ) is calculated as metric post-calibration minus metric pre-calibration. For the exact metric delta of each heatmap entry, see Table E1. **b)** Change in DE classification metrics post-sparsity calibration for cellular context generalization. For exact metric deltas see Table E2.

**Table E1:**
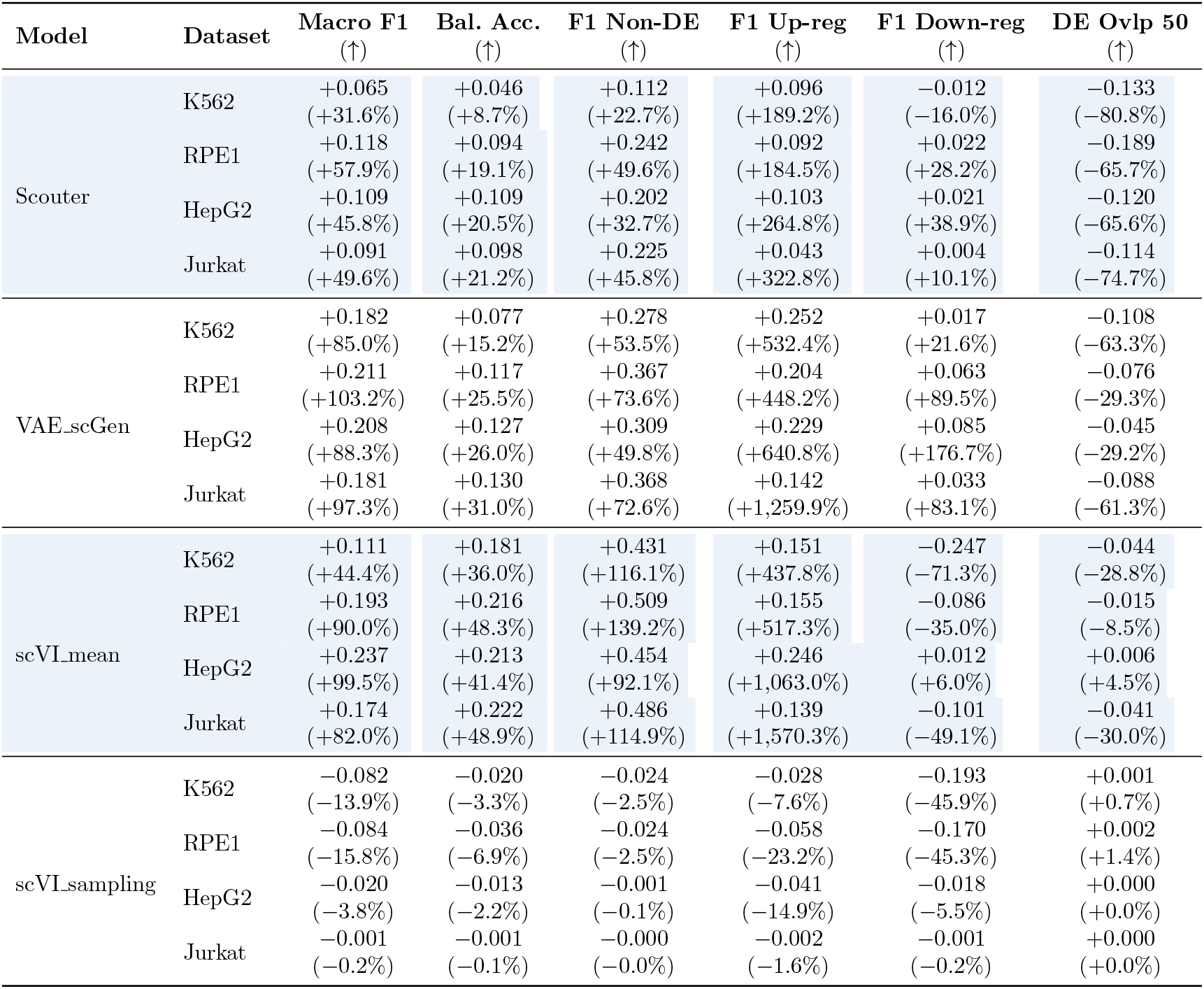
Values accompanying Figure E5a. Values report absolute difference (post - pre) with relative change in parentheses. ↑ denotes higher is better.

**Table E2:**
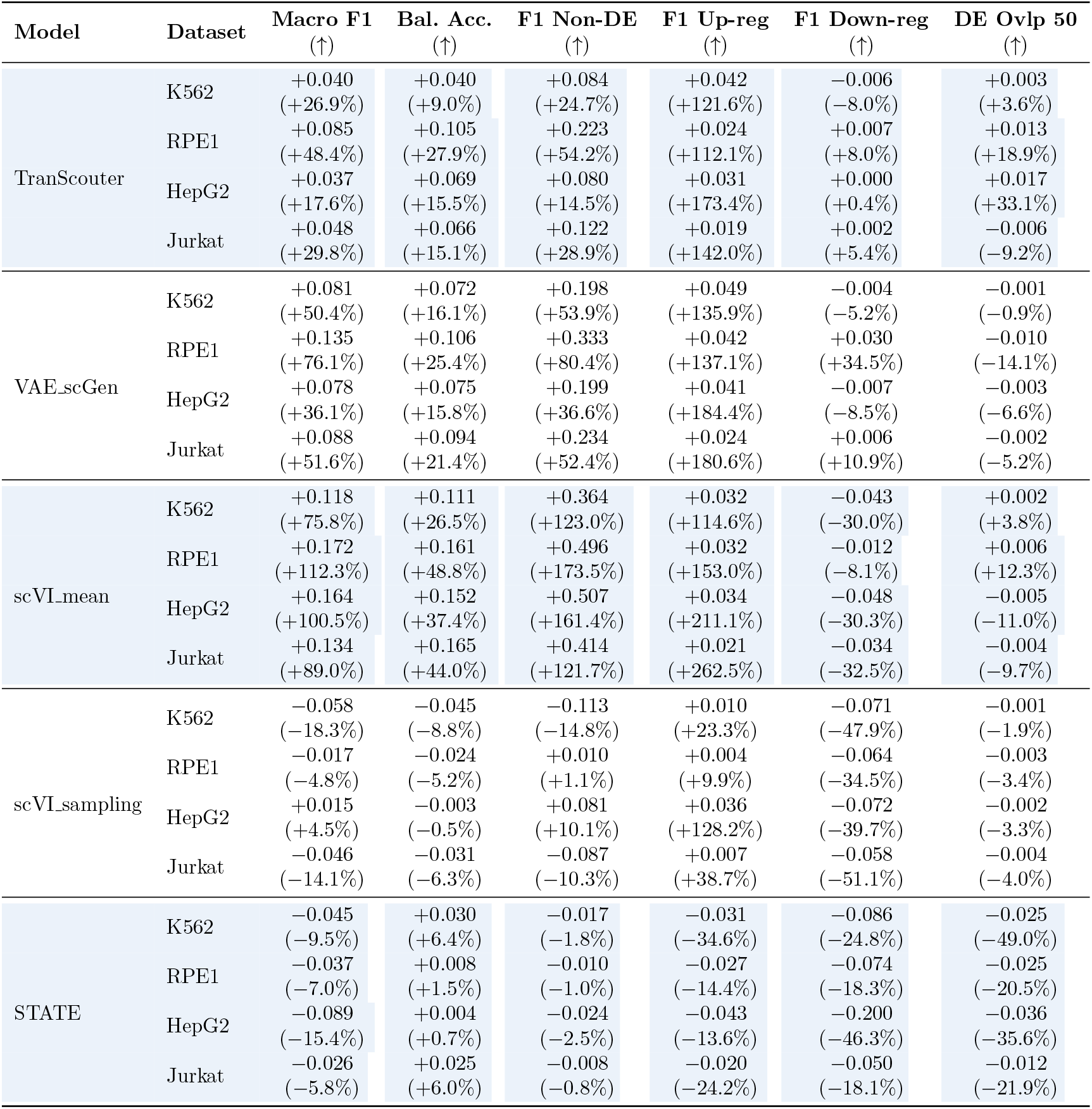
Values accompanying Figure E5b, right. Values report absolute difference (post - pre) with relative change in parentheses. ↑ denotes higher is better.

| Model | Dataset | Macro F1<br>(↑) | Bal. Acc.<br>(↑) | F1 Non-DE<br>(↑) | F1 Up-reg<br>(↑) | F1 Down-reg<br>(↑) | DE Ovp 50<br>(↑) |
| --- | --- | --- | --- | --- | --- | --- | --- |
| TranScouter | K562 | +0.040<br>(+26.9%) | +0.040<br>(+9.0%) | +0.084<br>(+24.7%) | +0.042<br>(+121.6%) | -0.006<br>(-8.0%) | +0.003<br>(+3.6%) |
|  | RPE1 | +0.085<br>(+48.4%) | +0.105<br>(+27.9%) | +0.223<br>(+54.2%) | +0.024<br>(+112.1%) | +0.007<br>(+8.0%) | +0.013<br>(+18.9%) |
|  | HepG2 | +0.037<br>(+17.6%) | +0.069<br>(+15.5%) | +0.080<br>(+14.5%) | +0.031<br>(+173.4%) | +0.000<br>(+0.4%) | +0.017<br>(+33.1%) |
|  | Jurkat | +0.048<br>(+29.8%) | +0.066<br>(+15.1%) | +0.122<br>(+28.9%) | +0.019<br>(+142.0%) | +0.002<br>(+5.4%) | -0.006<br>(-9.2%) |
| VAE_scGen | K562 | +0.081<br>(+50.4%) | +0.072<br>(+16.1%) | +0.198<br>(+53.9%) | +0.049<br>(+135.9%) | -0.004<br>(-5.2%) | -0.001<br>(-0.9%) |
|  | RPE1 | +0.135<br>(+76.1%) | +0.106<br>(+25.4%) | +0.333<br>(+80.4%) | +0.042<br>(+137.1%) | +0.030<br>(+34.5%) | -0.010<br>(-14.1%) |
|  | HepG2 | +0.078<br>(+36.1%) | +0.075<br>(+15.8%) | +0.199<br>(+36.6%) | +0.041<br>(+184.4%) | -0.007<br>(-8.5%) | -0.003<br>(-6.6%) |
|  | Jurkat | +0.088<br>(+51.6%) | +0.094<br>(+21.4%) | +0.234<br>(+52.4%) | +0.024<br>(+180.6%) | +0.006<br>(+10.9%) | -0.002<br>(-5.2%) |
| scVI_mean | K562 | +0.118<br>(+75.8%) | +0.111<br>(+26.5%) | +0.364<br>(+123.0%) | +0.032<br>(+114.6%) | -0.043<br>(-30.0%) | +0.002<br>(+3.8%) |
|  | RPE1 | +0.172<br>(+112.3%) | +0.161<br>(+48.8%) | +0.496<br>(+173.5%) | +0.032<br>(+153.0%) | -0.012<br>(-8.1%) | +0.006<br>(+12.3%) |
|  | HepG2 | +0.164<br>(+100.5%) | +0.152<br>(+37.4%) | +0.507<br>(+161.4%) | +0.034<br>(+211.1%) | -0.048<br>(-30.3%) | -0.005<br>(-11.0%) |
|  | Jurkat | +0.134<br>(+89.0%) | +0.165<br>(+44.0%) | +0.414<br>(+121.7%) | +0.021<br>(+262.5%) | -0.034<br>(-32.5%) | -0.004<br>(-9.7%) |
| scVI_sampling | K562 | -0.058<br>(-18.3%) | -0.045<br>(-8.8%) | -0.113<br>(-14.8%) | +0.010<br>(+23.3%) | -0.071<br>(-47.9%) | -0.001<br>(-1.9%) |
|  | RPE1 | -0.017<br>(-4.8%) | -0.024<br>(-5.2%) | +0.010<br>(+1.1%) | +0.004<br>(+9.9%) | -0.064<br>(-34.5%) | -0.003<br>(-3.4%) |
|  | HepG2 | +0.015<br>(+4.5%) | -0.003<br>(-0.5%) | +0.081<br>(+10.1%) | +0.036<br>(+128.2%) | -0.072<br>(-39.7%) | -0.002<br>(-3.3%) |
|  | Jurkat | -0.046<br>(-14.1%) | -0.031<br>(-6.3%) | -0.087<br>(-10.3%) | +0.007<br>(+38.7%) | -0.058<br>(-51.1%) | -0.004<br>(-4.0%) |
| STATE | K562 | -0.045<br>(-9.5%) | +0.030<br>(+6.4%) | -0.017<br>(-1.8%) | -0.031<br>(-34.6%) | -0.086<br>(-24.8%) | -0.025<br>(-49.0%) |
|  | RPE1 | -0.037<br>(-7.0%) | +0.008<br>(+1.5%) | -0.010<br>(-1.0%) | -0.027<br>(-14.4%) | -0.074<br>(-18.3%) | -0.025<br>(-20.5%) |
|  | HepG2 | -0.089<br>(-15.4%) | +0.004<br>(+0.7%) | -0.024<br>(-2.5%) | -0.043<br>(-13.6%) | -0.200<br>(-46.3%) | -0.036<br>(-35.6%) |
|  | Jurkat | -0.026<br>(-5.8%) | +0.025<br>(+6.0%) | -0.008<br>(-0.8%) | -0.020<br>(-24.2%) | -0.050<br>(-18.1%) | -0.012<br>(-21.9%) |

## Notes

### Competing Interest Statement

The authors have declared no competing interest.

